# Early frequency-specific contributions to serial-effects in audition

**DOI:** 10.1101/2023.01.22.525097

**Authors:** I. Lieder, A. Sulem, M. Ahissar

**Affiliations:** The Edmond and Lily Safra Center for Brain Sciences, The Hebrew University of Jerusalem, Jerusalem, Israel 9190401; Department of Psychology, The Hebrew University of Jerusalem, Jerusalem, Israel 9190401

## Abstract

Recent stimuli affect the perception of current stimuli, referred to as serial effects. These effects were mainly studied in the visual modality, where it was suggested that perceptual biases towards previous stimuli (contraction) stems from high-level processing stages, and promotes object-level stability. We now asked whether high object-level stages underlie contraction also in the auditory modality. We administered a two-tone pitch discrimination task using both pure and complex tones. Both have pitch, but they are perceived as different timbre categories. Pitch contraction was observed to be largest between tones of the same timbre-category, in line with the object-level account. To decipher the role of early, frequency-specific, category-indifferent processing-stages we used complex tones with missing fundamental. They differ in their low-level frequency components yet have the same pitch. Hence, a high-level account predicts that pitch contraction will remain. Surprisingly, we observed no contraction to the missing fundamental frequency. Rather, pitch was contracted to the physically-present frequencies. Supporting the low-level contribution, we found that though attention enhances contraction, it is not necessary. These observations suggest that contraction bias is an inherent part of the various stages of the auditory hierarchy of sensory processing.

## Introduction

Perception results from integrating responses to current sensory stimulation with prior expectations based on past experiences^1–3^. The integration with perceptual priors biases perception towards more expected stimuli and reduces the effect of external and internal noises^4–6^. *Serial dependence* (also called *serial effects*) refers to the effect of the most recent stimuli, including perceptual, decision and response biases^7,8^. These effects were most intensively studied in visual perception, where it was demonstrated for a broad range of simple (e.g., orientation^9–11^; position^12,13^) and complex features (e.g., facial identity^14,15^, facial gender and expression^16,17^, or facial attractiveness^18^). Serial effects were also found in other modalities, including olfactory^19^ and auditory stimuli^20–24^.

Although the behavioral signature of serial dependence has been widely investigated in the visual modality, its location along the hierarchy of sensory processing, perceptual decisions, and working memory is still debated. Most of the studies suggest that contraction to recent stimuli is formed either at high object-level processing stages and operates on global perceptual representations^10,13,14,18,25^, or at post-perceptual stages, involving decision making^26^ and working memory contributions^7,12,27,28,29^. The contribution of object-perceptual level is evidenced by the fact that the bias is manifested behaviorally even following trials with no requirement for a behavioral response, and that bias is apparent even when participants are asked to reply immediately, namely when no working memory is intentionally activated^13^. Differently, other observations such as that the magnitude of the bias on the perception of orientation^7^, position^12,27,28^ and facial emotional expression^29^ increases when a longer interval is imposed between perception and decision, suggest a contribution of working memory demands to serial effects. Additionally, serial dependence is found even between different types of stimuli, e.g., the orientation of Gabor stimuli biases orientation reproduction of dot patterns^30^. Both perceptual judgements and subsequent working-memory level accounts assume Bayesian optimization at object levels, which promotes perceptual likelihood and stability with respect to external objects and events.

On the other hand, a few visual studies suggested that the bias acts directly on sensory circuits. John-Saaltink et al.^31^ found that BOLD activity in the primary visual cortex (V1) is affected by the orientation of previous stimuli in the same position, even when no response is requested. However, the activity in V1 could arise from bottom-up contribution or rather results from feedbacks of higher levels on low-level representations^32^. Recently, Cicchini et al.^33^ suggested that priors are high-level constructs which integrate contextual information yet act upon early sensory signals, in line with the predictive coding framework.

These mixed results suggest two options. One, that different conditions give rise to biases at different processing stages, and hence different results are obtained in different protocols. Alternatively, the observed bias is the outcome of a hierarchy of biases, operating both at low- and at high-level processing stages. To distinguish between these options, we designed a protocol that allows quantitative assessment of biases at both low- and high-level processing stages within the same task in the auditory modality. We used the unique property of the relationship between frequency bands and the perceived pitch: along the fast-ascending pathway from the inner ear to the primary auditory cortex, processing is largely frequency specific, whereas later object-level stages are spectrally broad^34,35^. Thus, auditory objects, including simple ones, such as (harmonic) complex tones, are based on integration of the early spectrally-local frequency bands.

Complex tones are a superposition of the lowest fundamental frequency (first harmonic) and a series of harmonics (overtones that are integer multiples of the common fundamental frequency). Their perceived pitch is a high-level feature determined by the (sometimes inferred) fundamental frequency. Importantly, manipulating their spectral components (e.g., suppressing overtones or even the fundamental) while maintaining the periodicity of the acoustic signal, does not change their pitch. For example, complex sounds produced by different musical instruments, which share the fundamental frequency, have the same pitch, although having a different spectrum. This dissociation between the composing frequencies and the holistic feature indicates global, somewhat abstract pitch perception^36–41^. Thus, the components determine the high-level feature, but pitch perception is globally resilient to components. The relative intensities of the composing harmonics affect the perceived timbre, which is also a high-level feature, which is generated when the frequency bands are integrated. Thus, different musical instruments have different timbres, even when their pitch is shared.

Previous studies on serial effects in pitch perception have shown contraction to recent stimuli but have used pure tones, for which low-level and high-level processing stages might not be dissociated. Using complex tones, for which the local and global features are well differentiated, allowed us to dissect early from late perceptual biases in pitch perception. Having two timbre categories (pure and complex tones), we first tested whether pitch contraction is sensitive to timbre categories, namely whether contraction involves high-level influences of perceptual categories. Additionally, including complex tones, in particular with a missing harmonic (fundamental or second harmonic), allowed us to ask whether the effect also includes low-level representations, by testing whether contraction across timbre categories is to the low-level physically present frequency channels (harmonics), or to the high-level pitch feature. We applied computational tools (Generalized Additive Model, see Methods) that discriminated the effects of the recent (i.e., the previous trial) and long-term histories (i.e., all the previous trials in the session) to characterize the bias profile.

In a complementary study, we measured the contribution of top-down, task-driven versus bottom-up, stimulus-driven processes, to the magnitude of the contraction bias. We dissociated between these contributions by assessing the bias when performing the pitch discrimination task (with attention) versus a visually-presented arithmetic task while same tones are presented. Attention was shown to play an important role in contraction between visual stimuli^10^. But unlike our eyes, we cannot “close our ears”, and auditory attention, unlike vision, automatically tracks stimuli and deviations from regularities. This modality difference is manifested in the automatic event-related potential (ERP) response to auditory deviants (mismatch negativity^42^), while the visual mismatch negativity is more sensitive to participants’ attention^43^. We therefore reasoned that attention might not play a role in auditory contraction bias.

Together, these studies assessed two complementary aspects of the mechanisms underlying contraction bias: the contributions of low- and high-level processing stages and of bottom-up versus top-down processing directions.

### Study 1: The magnitude of serial effects is greater within than across timbre categories

We first asked whether complex tones contract pure tones and vice versa. These two types of tones belong to different categories of timbres, yet they have a shared feature – pitch. We measured contraction bias by the recent trial, across and within timbre category. We reasoned that if contraction is uniquely determined by abstract pitch – cross and within categories should be similar since they have the same abstract pitch (though different low-level representations). If contraction is determined uniquely by a low-level mechanism, bias magnitude will be determined by the distance between physical frequencies. If both abstract pitch (high-level) and frequency bands (low-level) contribute to contraction, we should find a larger within category, yet a significant cross category bias.

To test this, we administered a two-tone pitch discrimination task using both trials of pure and trials of complex tones, in random order (Methods: *Stimuli*). In each trial, two (220ms) consecutive tones (Stimulus 1, Stimulus 2) from the same timbre category (pure or complex), separated by a silent (800ms) inter-stimulus interval (ISI) were serially presented. Participants were asked to select which of the two tones had a higher pitch. We conducted three independent experiments. In Experiment 1, the complex tones were composed of three harmonics: fundamental frequency (*f*) and the first two overtones (*f*_1_ = 2 × *f*_0_ ; *f*_2_ = 3 × *f*_0_). In Experiment 2, the complex tones had a missing fundamental *f*_0_ and in Experiment 3, the complex tones had a missing second harmonic *f*_1_. Trials with pure tones and trials with complex tones were randomly intermixed, with equal probability. Thus, in each of the three experiments, there were four types of consecutive trials (with equal probability): two types in which consecutive trials had the same timbre (within-category trials, i.e., pure after pure and complex after complex) and two types in which consecutive trials had different timbres (across-category trials, i.e., complex after pure and pure after complex).

## Results

We evaluated the biases of pitch perception in each of the four types of consecutive trials: pure after pure, complex after complex, complex after pure and pure after complex (we merged the data of the three experiments; Methods: *Data analysis:* Study 1). Specifically, using GAM computational tools (see Methods), we assessed the profiles of the biases induced by recent history according to the similarity (in terms of frequency distance) between the current and previous stimuli. Since pitch contraction by recent occurs only within less than an octave^21,24^, we included only the (preceding) pure tone trials that were in the octave 500-1000Hz, which is the octave of the fundamental frequency of the complex tones (Methods: *Stimuli*: Studies 1 to 3). Figure 1a shows the magnitude of the biases as a function of the (log) frequency distance (*d*_1_) between the first tone in the current trial and the first tone in the previous trial. The shape of the curves is similar to that found by Lieder et al.^24^: contraction peaks at about half an octave and decreases with larger frequency distances. The two bias curves calculated for consecutive trials within-category, namely pure after pure and complex after complex, are largely overlapping. By contrast, the magnitudes of the cross-category biases are much smaller and are similar for both complex after simple and simple after complex trials.

**Figure 1.**
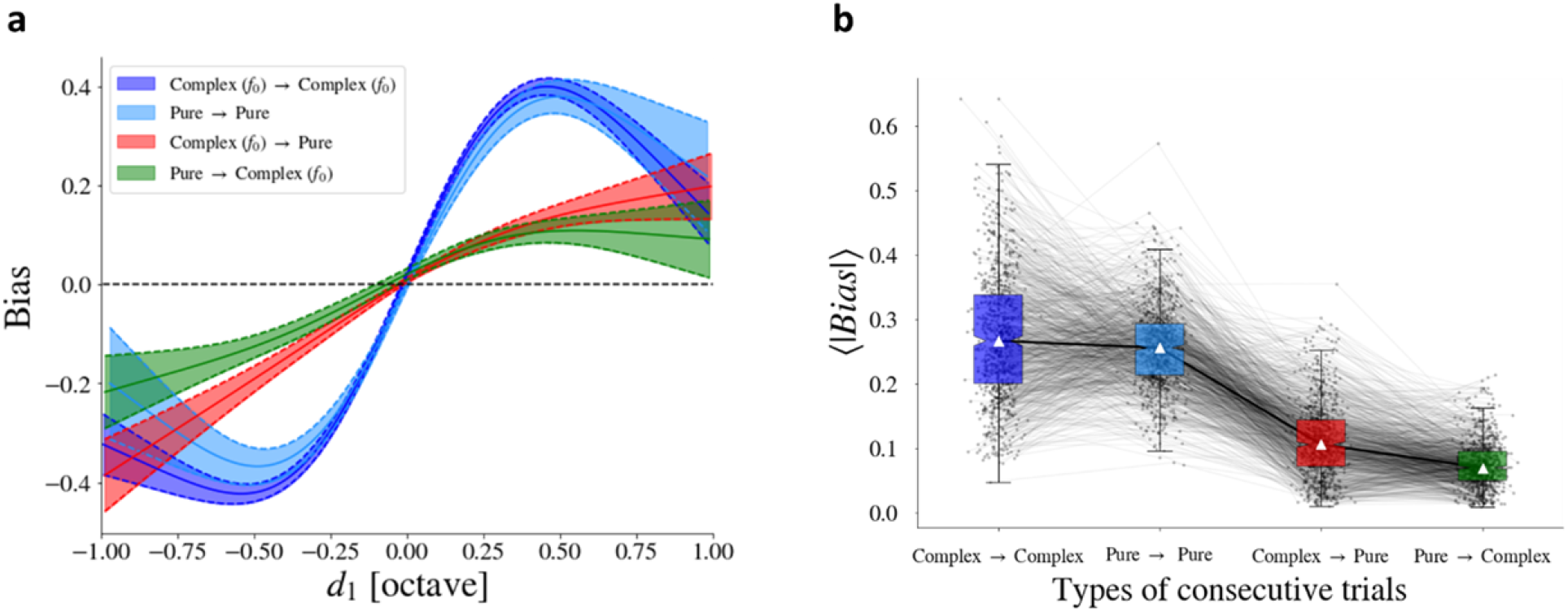
Serial effects are larger between trials of the same category than across timbre categories. Data from three experiments were aggregated in this analysis. **a**. The magnitude of the bias as a function of the log frequency distance (d_1_) between the first tone of current trial and the first tone of the previous trial. In the case of the harmonic complex tones, we calculated the log frequency distance from their fundamental frequencies. The fundamental frequencies of complex tones and frequencies of pure tones in preceding trials spanned 1 octave, 500-1000Hz, which is also the range spanned by the recent contraction bias. The magnitude of the bias is substantially larger between consecutive trials composed of the same timbre category (Complex *⟶* Complex, blue; Pure *⟶* Pure, cyan), than between consecutive trials composed of different timbre categories (Complex *⟶* Pure, red; Pure *⟶* Complex, green). Error bars indicate standard error. Black horizontal dashed line denotes zero bias. **b**. Mean of the absolute magnitude of individual participants’ bias in the four types of consecutive trials. White triangles indicate the median. Error bars show lower to upper quartile values of the data. Bias magnitudes are similar for consecutive trials within-category (Complex *⟶* Complex and Pure *⟶* Pure). Both are significantly higher than biases in across-category consecutive trials (Complex *⟶* Pure and Pure *⟶* Complex). Each line connects between the measures of the same participant under the four types of consecutive trial pairs.

This observation is also manifested in the comparison of the magnitude of the biases at the level of individual participants (Fig. 1b, Methods: *Data analysis*: Single participant fitting). The mean of the absolute value of the biases differed (*H* = 2107, *p* < .001, Kruskal-Wallis) between these conditions. Dunn’s test (with Bonferroni correction) showed that there was no difference between the two within-category biases (*Z* = 0.87, *p* = .99; Fig. 1b: blue compared to cyan boxplots), but each of them had a significantly larger bias than each of the two across-category biases (*Z* > 27, *p* < .001 for all comparisons; Fig. 1b: blue and cyan compared to red and green boxplots).

## Discussion

First, we replicated the shape and magnitude of the contraction curve found in Lieder et al.^24^ for consecutive trials of pure tones. In line with Lieder et al.^24^ and also with Chambers et al.^21^ (using complex Shepard tones), the contraction operates within a range of about one octave and peaks at about half an octave (tritone). It also corresponds to the general shape reported in the visual modality^10^. Calculating contraction between complex tones, we found an almost overlapping curve. However, the contraction across-categories is substantially smaller. Its shape seems different, but this may be an effect of insufficient statistics at the edges (larger frequency distances between consecutive trials), for which the number of trials is much smaller.

The substantially larger contraction within timbre categories suggests that contraction is contributed mainly by a late perceptual stage, after timbre categorization. This result is in line with the perceptual *continuity field* account in the visual modality^10^. According to this account, when there are no large obvious changes, high-level perception favors percepts that are similar to recent ones (several-seconds’ time window). For example, contraction between perceived facial expressions was found to be significantly stronger when faces of the same gender were presented in consecutive trials compared with faces of the opposite gender^44^. Similarly, Liberman et al.^16^ found that face-expression contraction is identity-specific. In a similar manner, in the auditory modality, sounds emitted by the same object (and consequently have the same timbre) manifest larger contraction bias. Such perceptual continuity helps us keep perceptual stability, in both visual and auditory modalities.

However, although substantially weaker, we also measured significant biases between consecutive trials belonging to different categories (complex after pure and pure after complex). This observation can be explained in two different ways. One, that serial effects across-category occur at an abstract level of pitch perception, though its magnitude is substantially smaller. According to this account, both within- and across-category biases operate at high level, and their magnitudes are enhanced within category. Second, that serial effects across-category precede timbre categorization, and occur before frequency channels are integrated into objects. The low and high levels share both the abstract high-level pitch and its concrete low-level component – harmonics. To determine between these accounts, we dissociated between the low-level frequency components and the perceived high-level pitch, by manipulating frequency bands while keeping their high-level features constant.

### Study 2: No Contraction when the Fundamental is missing

To determine whether serial effects across-categories operate globally, at the level of pitch perception, or locally, at the level of specific harmonics, we examined whether the physical presence of the fundamental frequency is necessary for contraction. Since the perceived pitch of complex tones is determined by the fundamental frequency, and that fundamental can still be perceived even when this frequency (or some overtones) is physically missing, a bias that operates only at the level of the perceived pitch should be insensitive to its physical presence. By contrast, if the bias operates at the level of separate frequency bands, complex tones with a missing fundamental might not contract, or not be contracted by, the perceived pitch.

## Results

### Pure after complex

First, we examined pure tone trials after complex tone trials under the three conditions: the full complex tones that included 3 harmonics (Experiment 1, Fig. 2a), complex tones with a missing fundamental (Experiment 2, Fig. 2b) and complex tones with a missing second harmonic (Experiment 3, Fig. 2c). For each condition, we concentrated on the contraction by the fundamental frequency (*f*_0_) on the frequency (*f*) of pure tones (within the same octave 500-1000Hz as *f*_0_ and hence within the contraction range, Methods: *Data analysis:* Studies 2 and 3). Namely, the contraction of *f* toward *f*_0_ (the pitch of the first pure tone perceived as closer to the pitch of the first complex tone in the previous trial). We calculated the magnitude of the biases as a function of the frequency distance (*d*_1_), between the fundamental frequency (*f*_0_) of the first complex tone in trial *t*−1, and the frequency (*f*) of the first pure tone in trial 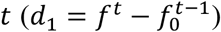 (Figs. 2d,e,f corresponding to Experiment 1, 2 and 3, respectively). We asked which representation of pitch the contraction operates on: the low-level representation (sensitive to the spectral composition of the stimuli) or the high-level representation (abstract and global) which operates even when *f*_0_ is missing.

**Figure 2.**
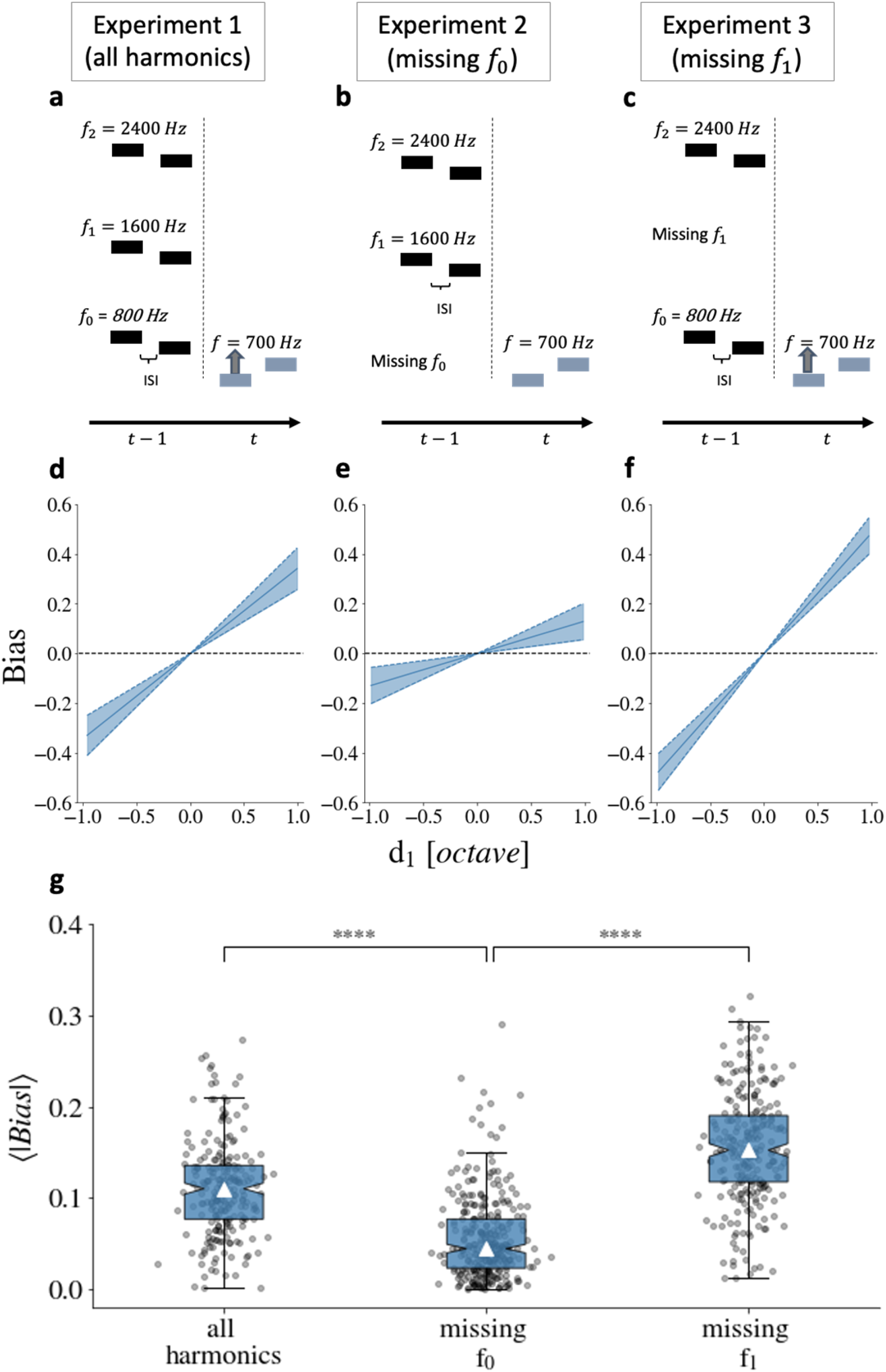
Contraction of a pure tone’s pitch by the perceived pitch of the preceding complex-tone trial (pure tone frequency within 500-1000Hz) when *f*_0_ is physically present (a, c, d, f) versus when it is missing (b, e) *−* there is no contraction by the pitch of the complex tone when its fundamental is missing. **a, b, c**. Schematic illustrations of complex tone trials at time *t*−1, followed by a pure tone trial at time *t*, under the three experimental conditions. **a**. Complex tones are composed of the three harmonics, *f*_0_, *f*_1_ and *f*_2_ (Experiment 1). **b**. Complex tones miss the fundamental and include only *f*_1_ and *f*_2_ (Experiment 2). **c**. Complex tones are composed of *f*_0_ and *f*_1_ and miss *f*_1_ (Experiment 3). **d, e, f**. Contraction biases measured as a function of the frequency distance 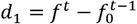. Error bars indicate standard error, based on the Bayesian posterior distribution of the bias slope. Black horizontal dashed line denotes zero bias. **g**. Comparison of the bias magnitudes within individual participants when: left - all frequencies are physically present; middle – *f*_0_ is missing; and right – *f*_1_ is missing. The bias magnitude was calculated for each participant using as a metric the mean of the absolute value of the bias magnitude. White triangles indicate the median. Error bars show lower to upper quartile values of the data. Most participants showed contraction (the slopes of the bias function were significantly positive): 227/230 participants in Experiment 1 (*w* = 11, *p* < .001, Wilcoxon signed-rank), 250/253 in Experiment 3 (*w* = 46, *p* < .001, Wilcoxon signed-rank), and 254/316 when *f*_0_ was missing (Experiment 2), (*w* = 6583, *p* < .001, Wilcoxon signed-rank). The analysis of both aggregate (**d, e, f**) and individual data (**g**) showed that serial effects were substantially and significantly reduced when pure-tone trials followed complex-tone trials with a missing fundamental, suggesting that serial effects occurred through frequency channels. **** *p* < .001.

To test it, we compared the contractions when the fundamental frequency was physically present (Experiments 1 and 3) to its magnitude when the fundamental frequency was missing (Experiment 2). When the fundamental was missing, the slope of the bias was almost flat (Fig. 2e), *slope* = 0.13 ± 0.08 (SE), and non-significantly different from zero (*p* = 0.095). Namely, there was no contraction by the pitch when the fundamental was missing. By contrast, when *f*_0_ was physically present, the slopes were substantially steeper and highly significant: *slope* = 0.37 ± 0.09 (SE), *p* < .001 (Experiment 1, Fig. 2d); *slope* =0.53 ± 0.08 (SE), *p* < .001 (Experiment 3, Fig. 2f). The predictor *d*_1_ had a highly significant contribution to the model (*χ*^2^ = 125, *edf* = 78, *p* = .0005; *χ*^2^ 189, *edf* = 103, *p* < .001, Wald test; Experiment 1 and Experiment 3, respectively), though it was also significant when *f*_0_ was missing (*χ*^2^ = 159, *edf* = 113, *p* = .003, Wald test; Experiment 2), and contraction was very small and close to zero.

These results were also evident at the level of individual participants (Fig. 2g plot the mean of the absolute value of the magnitude of biases, Methods: *Data analysis*: Single participant fitting). Though most participants showed contraction (the slopes of the bias were significantly positive) in all three experiments, the magnitude of the effect (calculated as the mean of the absolute value of individual participants’ bias magnitudes) differed between these conditions (*H* = 337.5, *p* < .001, Kruskal-Wallis). Post-Hoc tests (Bonferroni correction, Dunn’s test) showed that the bias was larger when *f*_0_, *f*_1_ and *f*_2_ were physically present (Experiment 1) compared to when *f*_0_ was missing (Experiment 2): (*Z* = 10.8, *p* < .001). Similarly, the bias was significantly larger when *f*_0_ and *f*_2_ were physically present (Experiment 3) compared to when *f*_0_ was missing (Experiment 2): (*Z* = 18.1, *p* < .001).

### Complex after pure

We now consider the reverse order of trials, i.e., complex-tone trials after pure-tone trials, under the three conditions: the full complex tones (Experiment 1, Fig. 3a), missing fundamental (Experiment 2, Fig. 3b) and missing second harmonic (Experiment 3, Fig. 3c). For each condition, we measure the contraction that is driven by the first pure tone of the previous trial (*f*^*t*−1^) and acts on the fundamental frequency of the first complex tone of the current trial 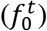. This bias indicates that the pitch of the complex tone was perceived as closer to the pitch of the pure tone. As mentioned above, pure tones were in the contraction range of *f*_0_ as they were in the same octave 500-1000Hz (Methods: *Data analysis:* Studies 2 and 3). We calculated the magnitude of the biases as a function of the log frequency distance (*d*_1_) between the preceding pure tone (*f*^*t*−1^) and the current fundamental frequency 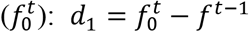 (Figs. 3d,e,f, corresponding to Experiment 1, 2 and 3, respectively). Using the same rationale as in the previous analysis, we examined whether it is necessary that the fundamental frequency will be present for contraction to occur.

**Figure 3.**
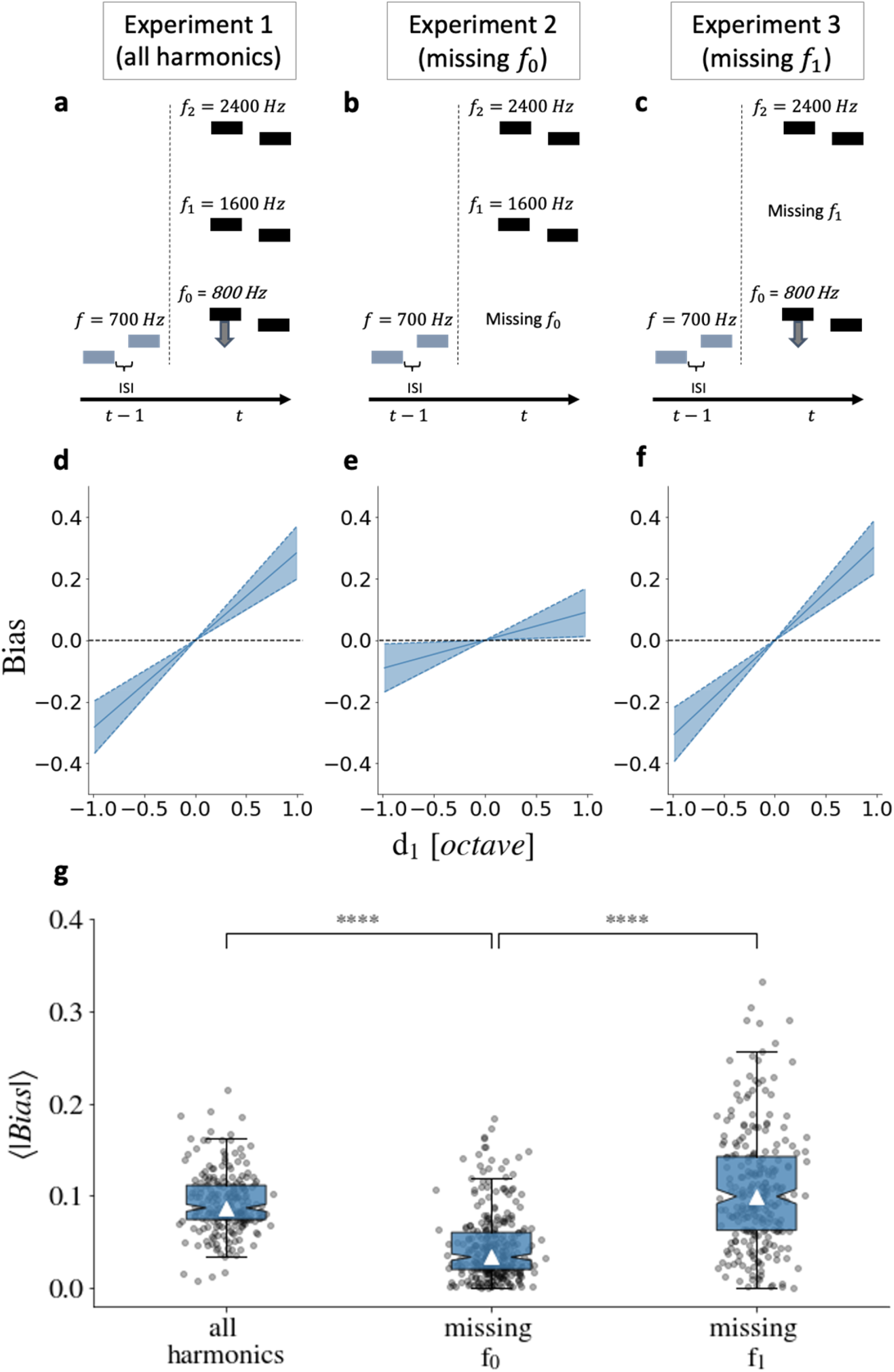
Contraction of the perceived pitch of a complex tone by the preceding pure-tone trial (pure tone frequency within 500-1000Hz) where *f*_0_ is physically present (a, d, c, f) versus when it is missing (b, e) *−* there is no contraction of the pitch of the complex tone when the fundamental is missing. **a, b, c**. Schematic illustrations of pure-tone trials at time *t*−1, followed by a complex-tone trial at time *t*. **a**. Complex tones are composed of the three harmonics, *f*_0_, *f*_1_ and *f*_2_ (Experiment 1). **b**. Complex tones include only *f*_1_ and *f*_2_, and miss *f*_0_ (Experiment 2). **c**. Complex tones include only *f*_0_ and *f*_2_, and miss *f*_1_ (Experiment 3). **d, e, f**. Contraction biases measured as a function of the frequency distance 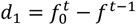. Error bars indicate standard error. Black horizontal dashed line denotes zero bias. **g**. Comparison of the bias magnitudes within individual participants when: left – all frequencies are physically present; middle – *f*_0_ is missing; and right – *f*_2_ is missing. The bias magnitude was calculated for each participant using as a metric the mean of the absolute value of the bias’s magnitude. White triangles indicate the median. Error bars show lower to upper quartile values of the data. Most participants showed contraction (the slopes of the bias function were significantly positive): 227/230 participants in Experiment 1 (W=8; p<.001, Wilcoxon signed-rank), 232/253 in Experiment 3 (W=942; p<.001, Wilcoxon signed-rank), and 239/316 participants when *f* was missing (Experiment 2), (W=9288, p<.001, Wilcoxon signed-rank). The analysis of both aggregate (**d, e, f**) and individual data (**g**) showed that the pitch of a complex tone is not contracted toward the preceding pure tone when the fundamental is missing, thus challenging the account that the effect across-categories acts on global pitch perception. Furthermore, when the fundamental is physically present, the contraction of pitch of complex tones is substantially and significantly stronger.

We compared the contractions when the fundamental frequency was physically present (Experiments 1 and 3) to its magnitude when the fundamental frequency was missing (Experiment 2). In line with the above result, in the missing fundamental condition (Experiment 2), the slope of the bias was flat (Fig. 3e), *slope* = 0.07 ± 0.07 (SE) and not significantly different from zero (*p* = .32). Namely, there was no contraction of the pitch when the fundamental was missing. By contrast, when the fundamental was physically present, the slopes of the bias were substantially steeper and highly significant: *slope* = 0.26 ± 0.08 (SE), *p* < .001 (Experiment 1, Fig. 3d); *slope* = 0.33 ± 0.08 (SE), *p* < .001 (Experiment 3, Fig. 3f), and the predictor *d*_1_ had significant contribution to the model (*χ*^2^ = 58.5, *edf* = 41, *p* = .04; *χ*^2^ = 569.0, *edf* = 215, *p* < .001, Wald test; Experiment 1 and Experiment 3, respectively), but not when *f*_0_ was missing (*χ*^2^ = 53.2, *edf* = 76, *p* .98, Wald test; Experiment 2).

Analyses at the level of individual participants yielded the same results (Fig. 3g plot the mean of the absolute value of the participants’ bias magnitude, Methods: *Data analysis*: Single participant fitting). Though most participants showed contraction (the slopes were significantly positive) in all three experiments, the magnitude of the bias differed between these conditions (*H* = 246.8, *p* < .001, Kruskal-Wallis). Post-Hoc tests (Bonferroni correction, Dunn’s test) showed that bias was larger when *f*_0_, *f*_1_ and *f*_2_ were physically present (Experiment 1) compared to when *f*_0_ was missing (Experiment 2): (*Z* = 12.9, *p* <). Similarly, the bias was significantly larger when *f*_1_ and *f*_2_ were physically present (Experiment 3) compared to when *f* was missing (Experiment 2): (*Z* = 13.7, *p* < .001).

## Discussion

We observed no contraction across-category trials when the fundamental was missing. Furthermore, the contraction was substantially stronger and highly significant when the fundamental was physically present. By dissociating the low-level from the high-level representations, we demonstrated that, across-category, there is no contraction to the (global) pitch but to the specific frequency *f*_0_ when physically present. Together, these results imply that low-level contraction is manifested in cross-category trials.

### Study 3: Contraction to physical harmonics that differ from the perceived pitch

In this study we asked whether the physical presence of a frequency band is sufficient for contraction. We reasoned that if contraction operates early, the physical presence of a frequency will induce contraction even when it differs from the globally-perceived pitch. We focused on pure-tone trials within the upper octave (1000-2000Hz) that followed a complex-tone trial. Importantly, the frequency (*f*) of the pure tones was in the same octave as the second harmonic (*f*_1_) of the complex tones and thus, within their contraction range (Methods: *Stimuli*: Studies 1-3). We measured the contraction of *f* of the first pure tone in the current trial by *f*_1_ of the first complex tone in the preceding trial. We compared the bias magnitude when *f*_1_ was physically present (Experiments 1 and 2) to when *f*_1_ was missing (Experiment 3). Crucially, unlike *f*_0_, *f*_1_ does differ from the perceived pitch. Thus, measuring contraction of pure tone by *f*_1_ when the latter is physically present would provide strong evidence for a low-level bottom-up account, namely that the bias operates at an early processing stage, via feedforward pathways.

Note that the bias in the reverse order of trials, namely the bias driven by the first pure tone of the preceding trial (*f*^*t*−1^ – in the high 1000-2000Hz octave) on the second harmonic of the first complex tone of the current trial 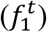, cannot be measured. Our analysis is based on participants’ pitch judgements. However, in the case of complex-after pure-tone trials, the response of the current trial addresses the pitch of the complex tones, which is associated with *f*_0_ and not with *f*_1_ .

## Results

We analyzed pure tone trials following complex tone trials, when pure tones were within the same octave (1000-2000Hz) as the second harmonic (*f*_1_) (Methods: *Data analysis:* Studies 2 and 3), under the three conditions: the full complex tones that included 3 harmonics (Experiment 1, Fig. 4a), complex tones with a missing fundamental (Experiment 2, Fig. 4b) and complex tones with a missing second harmonic (Experiment 3, Fig. 4c). For each condition, we calculated the magnitude of the bias as a function of the frequency distance 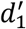 to the second harmonic 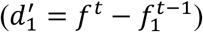, namely the contraction of *f* to *f*_1_ (Figs. 4d,e,f corresponding to Experiment 1, 2 and 3, respectively). The bottom-up frequency-channel account predicts that contraction of pure tones by *f*_1_ will occur only when the latter is physically present, even though different from pitch.

**Figure 4:**
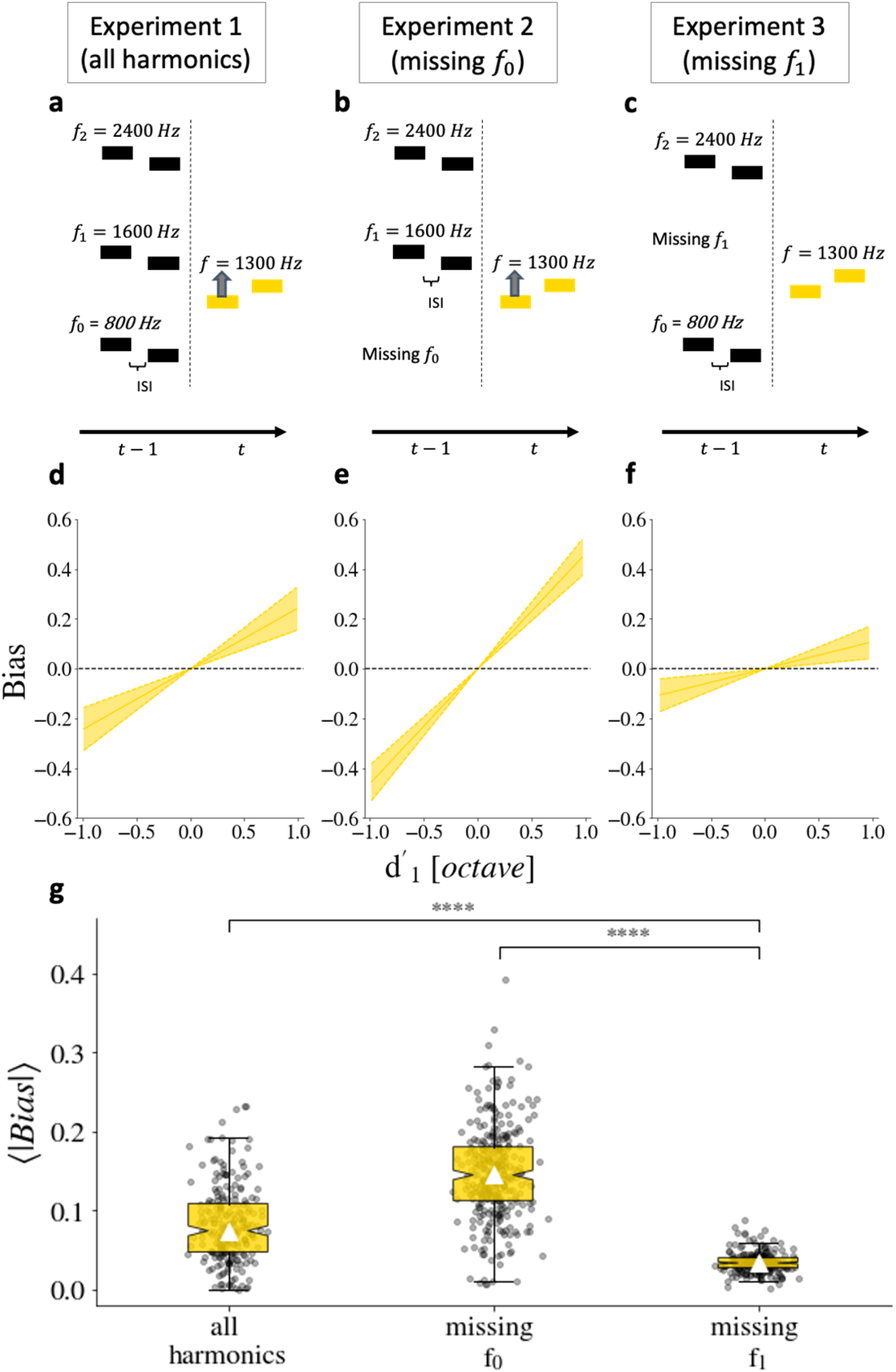
Contraction of a pure-tone pitch (within 1000-2000Hz) by the second harmonic (*f*_1_ - in the same frequency range) of a preceding complex-tone trial when *f*_1_ is present (a, b, d, e) versus when it is missing (c, f) *−* pure tones are contracted by *f*_1_ when present, even though it differs from the perceived pitch (determined by *f*_0_). **a**,**b**,**c**. Schematic illustrations of complex tone trials at time *t*−1, followed by a pure tone trial at time *t* in the same octave as *f*_1_. **a**. Complex tones are composed of the three harmonics, *f*_0_, *f*_1_ and *f*_2_ (Experiment 1). **b**. Complex tones include only *f*_1_ and *f*_2_, and miss *f*_1_ (Experiment 2). **c**. Complex tones include only *f*_0_ and *f*_2_, and miss *f*_1_ (Experiment 3). **d, e, f**. Contraction biases measured as a function of the frequency distance 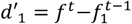. Error bars indicate standard error, based on the Bayesian posterior distribution of the bias slope (*w*′_1_). Black horizontal dashed line denotes zero bias. **g**. Comparison of the bias magnitudes within individual participants when: left – all 3 harmonics are physically present; middle – *f*_0_ is missing; and right – *f*_1_ is missing. The bias magnitude was calculated for each participant using as a metric the mean of the absolute value of the bias’s magnitude. White triangles indicate the median. Error bars show lower to upper quartile values of the data. As in the previous analysis, most participants showed contraction (the slopes of the bias function were significantly positive): 209/230 participants in Experiment 1 (*w* = 853, *p* < .001, Wilcoxon signed-rank), 313/316 in Experiment 2 (*w* = 14, *p* < .001, Wilcoxon signed-rank), and 252/253 in Experiment 3 (*w* = 3.0, *p* < .001, Wilcoxon signed-rank), although bias magnitude was very low when *f*_1_ was missing.

We thus compared the contractions when *f*_1_ was present (Experiments 1 and 2) to its magnitude when *f*_1_ was missing (Experiment 3). When *f*_1_ was missing, the slope of the bias was very shallow (Fig. 4f) and not significant: *slope* = 0.08 ± 0.07 (SE), *p* = .25; namely, there was no contraction by *f*_1_ when it was missing. By contrast, when *f*_1_ was present, the contraction was much stronger and significant: *slope* = 0.29 ± 0.09 (SE), *p* = .0017 (Experiment 1, Fig. 4d); *slope* = 0.48 ± 0.08 (SE), *p* < .001 (Experiment 2, Fig. 4e). Furthermore, the predictor 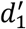 had a significant contribution to the model when *f*_1_ was present (*χ*^2^ = 127, *edf* = 84, *p* = .0016; *χ*^2^ = 234, *edf* = 132, *p* < .001, Wald test; Experiments 1 and 2, respectively), but not when *f*_1_ was missing (*χ*^2^ 35, *edf* = 31, *p* = .32, Wald test; Experiment 3).

Statistical analyses at the level of individual participants show the same results (Fig. 4g display the mean of absolute value of the biases, Methods: *Data analysis*: Single participant fitting). As in the previous analyses, most participants showed contraction (the slopes were significantly positive) in all three experiments but the magnitude of the biases differed between these conditions (*H* = 463.5, *p* < .001, Kruskal-Wallis). Post-Hoc tests (Bonferroni correction, Dunn’s test) showed that the bias was larger when *f*_0_, *f*_1_ and *f*_2_ were present (Experiment 1) compared to when *f*_1_ was missing (Experiment 3): (*Z* = 10.4, *p* < .001). Similarly, the bias was significantly larger when *f*_1_ and *f*_2_ were physically present (Experiment 2) compared to when *f*_1_ was missing (Experiment 3) missing: (*Z* = 21.5, *p* < .001).

## Discussion

We found significant contraction, driven by a physically present harmonic which is not associated with the perceived pitch, on pure tone within the one-octave contraction range. Thus, the existence of a harmonic is sufficient for contraction. This finding indicates that contraction operates locally at the level of specific frequency bands with energy, in a bottom-up manner. Had it been a top-down effect operating at a low frequency-specific level, it would have yielded contraction consistent with the globally perceived pitch. This result is also consistent with Chambers et al.^21^, who explored the influence of context effects on the perception of ambiguous stimuli and proposed a computational model for contraction, which operates at the level of frequency channels, and supports spectro-temporal continuity of sound sources. Temporal binding of successive frequency components, operating at early processing stages, before conscious perception, may also contribute to stabilize perceptual organization and to reduce noise. Together, Studies 2 and 3 show that in cross-category contraction, low-level physical proximity of the physically present frequency is both necessary (Study 2) and sufficient (Study 3), supporting a low-level bottom-up contribution to perceptual contraction.

### Study 4: Attention enhances contraction bias

In Study 4, we administered an experimental protocol which enabled us to measure the impact of task-related attention on the magnitude of bias by the recent trial. For that, participants were asked to perform one of two tasks in each trial, interchangeably. In half of the trials (randomly selected) participants were asked to perform the two-tone pitch discrimination task (using pure tones), and in the other half, participants were asked to perform a simple arithmetic task. Two pure tones, with same temporal structure were presented in all trials. Hence, participants were exposed to the same 2-tone statistics in all trials, and only the attentional mode differed. This structure allowed us to compare contraction in trials that were preceded by the pitch discrimination task to that in trials that were preceded by simple 1-digit multiplication confirmation, and thereby quantify the contribution of attention.

Each trial began with a visual script (500ms): “sounds” or “math” which indicated the required task in that trial (Figure 5). In both cases, two (220ms) consecutive pure tones (Stimulus 1, Stimulus 2) separated by a silent (800ms) inter-stimulus interval (ISI) were presented. In “sounds” trials, participants were asked to select which of the two tones has a higher pitch. In “math” trials, two multiplications were presented visually, temporally aligned with the two tones, and participants were asked to determine which of the two products is greater. In this case, successful task performance requires attention to the arithmetic task. Thus, one half of the “sounds” trials followed a trial with selective attention to the pitch (“sounds” trial), and the other half followed a trial (“math” trial) with selective attention to the math task. The mean accuracy of correct responses was 89 ± 13.8% (SD) for the “math” task and 77.8 ± 14.3% (SD) for the “sounds” task.

**Figure 5.**
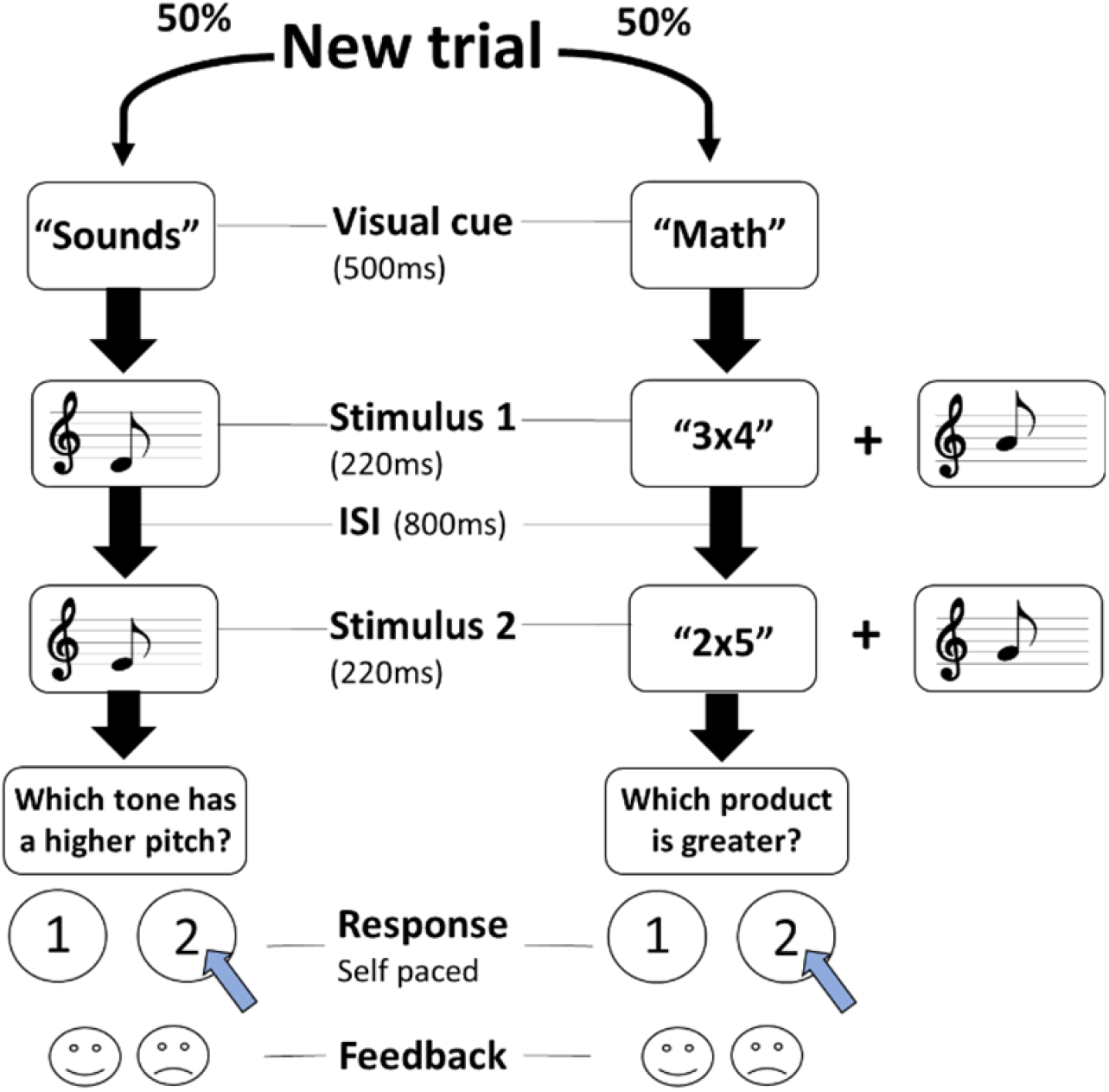
Illustration of the temporal structure of the two trial types. Left – 2-tone discrimination. Right – simple arithmetic comparisons. Each trial began with one of the two possible visual cues, presented for 500ms, “sounds” (in which attention is focused on the auditory task) or “math” (in which attention is focused on arithmetic comparison task), randomly chosen with equal probability. In both trial types, two consecutive pure tones (Stimulus 1, Stimulus 2), separated by a silent inter-stimulus interval (ISI) were presented. In “sounds” trials, following the tone presentations, participants were asked to determine which tone had a higher pitch by pressing “1” or “2”. In “math” trials, a simple multiplication of digits from 1 to 5 is presented, with each tone, and participants were asked to determine which of the two products was larger. Visual feedback followed each response.

## Results

We compared the magnitude of pitch contraction in trials that followed “math” trials to pitch contraction in trials that followed “sounds” trials. Figure 6a displays the bias as a function of the frequency distance between the first tone in the current trial and the first tone in the previous trial 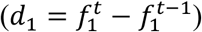. The shape of the contraction function was similar in both cases – contraction peaks at half an octave away (tritone), and decreases with larger frequency distances. Yet, when attention was directed toward the tones, the magnitude of the bias was substantially larger. Comparing the bias under these attentional states within individual participants (Methods: *Data analysis*: Fitting single participants; Fig. 6b), showed a highly significant difference (*U* = 12702, *p* < .001, Mann-Whitney U test).

**Figure 6.**
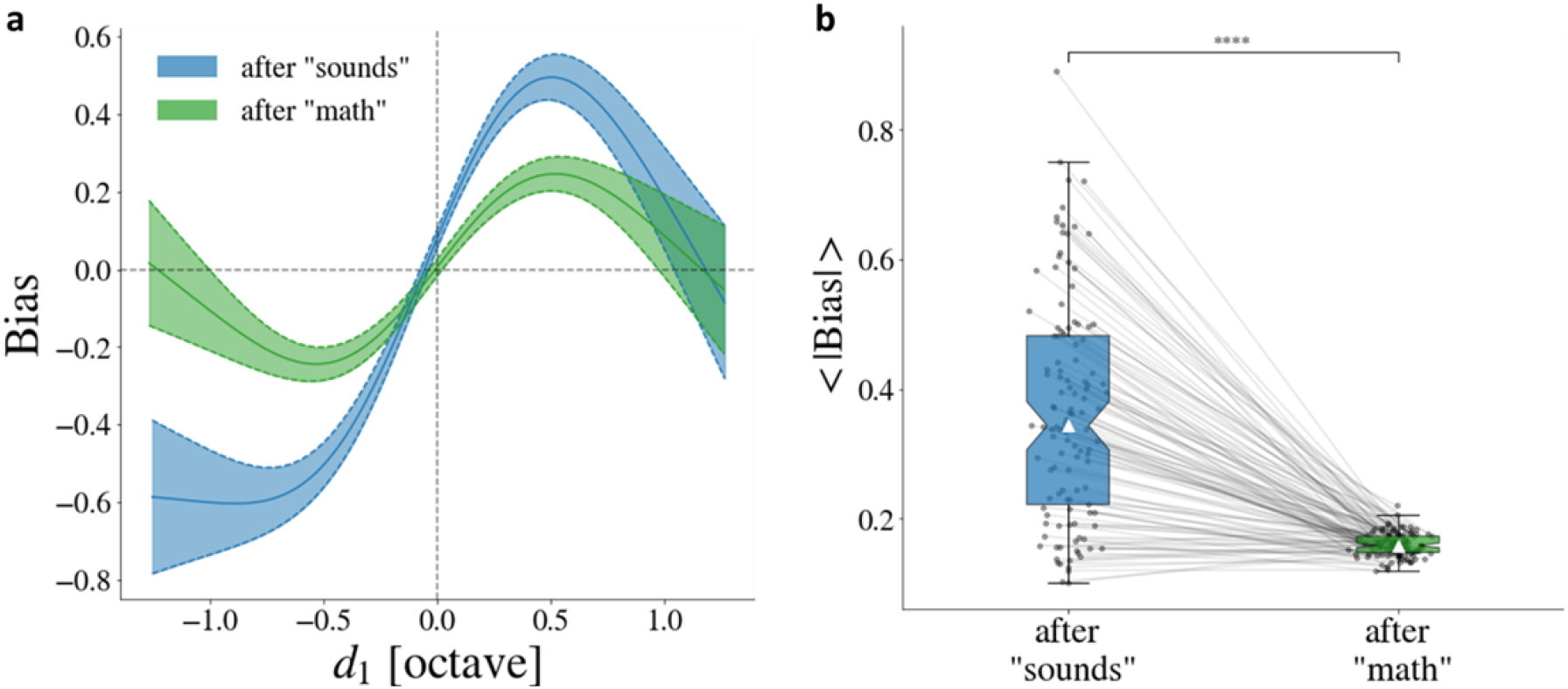
The effect of attention: contraction is larger after “sounds” trial, in which participants direct their attention to the pitch task, than after “math” trials, in which their attention was actively directed toward the arithmetic comparison task. **a**. The magnitude of the bias as a function of the frequency distance d_1_ between the first tone of the current and the previous trials. Though both curves have the same shape, and both peak at ∼0.5 octave, the magnitude of the bias is substantially larger with task-related attention in previous trials. Error bars indicate standard error. Black horizontal and vertical dashed lines denote zero bias and zero frequency distance, respectively. **b**. Magnitude of the biases of individual participants in trials following “sounds” and trials following “math”. Each line connects between bias magnitudes calculated for the same participant under the two attentional states. White triangles indicate the median. Error bars show lower to upper quartile values of the data. **** *p* < .001.

## Discussion

We found that following trials in which attention was actively directed toward the pitch discrimination task, the bias magnitude was substantially enhanced compared to following trials in which attention was directed to the arithmetic comparison task. The shape of the function was similar in both cases and was consistent with that previously reported for tone pitch discrimination^24^. The observation that serial effects rely on attention implies that its underlying mechanism involves later, attention demanding, stages along the perceptual processing^45^. However, although weaker, there is also contraction when attention in the previous trial was actively directed to an alternative task.

Our findings on the contribution of attention in the auditory modality are in line with observations made in the visual modality. Fischer & Whitney^10^ demonstrated that the strength of serial dependence in Gabor orientations is enhanced by spatial attention in the previous trial. Stronger bias occurs for stimuli for which the cued location remains constant from the previous to the current trial. Fritsche & de Lange^46^ showed that the magnitude of serial dependence is affected by the attended feature. Specifically, they found that serial dependence of orientation was markedly reduced when observers attended to the size, rather than the orientation of the previous trial.

While attention has a large effect on the magnitude of the serial dependence, the actual motor response, is probably not crucial. Previous studies found a similar magnitude of contraction even when a motor response was not required in the previous trial^10,13^ and even in the absence of any explicit task^47^. Our experimental protocol is slightly different, in that we asked the participants to perform an arithmetic discrimination task instead of the pitch discrimination task, at the time they are exposed to the auditory stimuli. Thus, attention was actively directed toward another task.

The substantial contribution of attention to contraction bias differs from its minor to no contribution to mismatch responses to pitch deviants^48^. This difference is surprising, since mismatch negativity is also based on (automatic) tracking of previous stimuli, and forming implicit predictions regarding subsequent stimuli. When these are surprising and do not match predictions, the response is enhanced. Perhaps mismatch negativity shares an earlier stage with contraction bias, but contraction also has an additional higher-level component, as evident in Study 1.

### General discussion

In this study, we used advanced computational tools to investigate perceptual bias of pitch, induced by recent stimuli. We focused on the contributions of low- and high-level processing stages and on their processing dynamics, dissociating bottom-up from attention driven contributions. By mixing trials of different timbre categories (pure and complex tones), we showed that serial effects in pitch are substantially stronger between stimuli of the same category, suggesting a contribution of high, object-level processing stages to the bias magnitude. This result is in line with previous studies in the visual modality, supporting the suggestion of a *continuity field* at the level of task-relevant objects^16,18,49^. Recently, Manassi & Whitney^50^ demonstrated that serial effects can cause an illusion of visual stability in a continuously changing object. Furthermore, several studies have shown a contribution of higher, post-perceptual stages, including inertia in decision-making^26^ and working memory biases^7^. For example, serial effects on abstract representations of orientation have been shown between different types of stimuli that share abstract orientation but not its visual representation^30^.

In Studies 2 and 3, we characterized the contraction operating between trials of different timbre categories. Surprisingly, we found contraction bias within physically present frequency bands, implying an early, pre-frequency-convergence component. This component does not act on the abstract frequency-global representation of pitch, but on low-level stimulus representations. Namely, it is not driven by the percept, suggesting the contribution of bottom-up, stimulus-driven processes.

In Study 4, we examined the contribution of bottom-up and top-down contributions by manipulating selective attention. Spatial attention has a major contribution in the visual modality^10^, but, since the auditory modality tracks stimuli statistics more automatically, as manifested in the mismatch negativity phenomenon^43^, the role of attention in the auditory modality could be reduced. We found that also in the auditory modality, attention substantially enhanced bias magnitude, indicating top-down contributions. Still, a significant bias was also observed when attention was actively directed to another task, implying bottom-up contributions.

Together, the results of these four studies demonstrate that the measured serial contraction bias is the combined outcome of both low-level and high-level representations and is contributed by both bottom-up (stimulus-driven) and top-down (task-driven) projections. This complex pattern suggests that context biases may be an integral part of each stage along the processing hierarchy^51^, implemented by neural retention processes, such as adaptation^52,53^, which enable familiarity and predictions of environmental statistics at its various spectral and temporal scales.

## Methods

### Participants

The experiment was conducted via the Amazon Mechanical Turk (M-Turk) platform. All participants (all from the United States) were recruited with previous M-Turk approval >95% and a total number of human intelligence tasks (HITs) >1,000. Our written instructions emphasized that the experiment must be performed: (1) using headphones in a quiet environment; (2) with either a laptop or a desktop computer; and (3) only by people with good hearing who are between the ages of 20 and 50. Each individual could only participate once. In all the experiments, participants performed 300 trials (three blocks of 100 trials each) that lasted between 10 to 15 minutes and were paid US$2. In all the Studies, we excluded participants whose accuracy ≤ 55% correct, since this low accuracy suggests that they may not have been performing the task. This yielded 22.9%, 17% and 19.5% exclusion in Experiments 1, 2, 3 (Studies 1-3), respectively and 8.5% exclusion in Study 4. The mildly high rate of exclusion is largely an outcome of our choice of a non-adaptive protocol, which is more challenging, but ensures that same stimuli statistics is used for all participants. We also assessed the consistency of performance throughout the task, by measuring the variability of the mean accuracy in windows of 60 trials. This additional criterion led to 2.9%, 4% and 5.6% exclusions in Experiments 1, 2, 3 (Studies 1-3), respectively and did not lead to further exclusions in Study 4.

#### Studies 1-3

We recruited 310 (Experiment 1), 400 (Experiment 2) and 338 participants (Experiment 3).

– In Experiment 1, their average age was 34.7 years (*SD* = 7.9), 46.1% were females, and they had in average 2.3 years of musical experience (*SD* = 4.0).
– In Experiment 2, their average age was 34.0 years (*SD* = 7.6), 44.8% were females, and they had in average 2.4 years of musical experience (*SD* = 4.6).
– In Experiment 3, their average age was 34.8 years (*SD* = 8.2), 45.0% were females, and they had in average 2.6 years of musical experience (*SD* = 4.5).

The data of 230 (74.2%) (Experiment 1), 316 (79.0%) (Experiment 2) and 253 (74.9%) (Experiment 3) participants are included.

#### Study 4

We recruited 153 participants. They were asked to give their identifying M-Turk worker ID, age (*M* = 31.8 years, *SD* = 5.4), gender (41.8% females) and musical experience (*M* = 2.6 years, *SD* = 4.4). In addition to the excluded participants whose accuracy in the “sounds” task was ≤55% correct (13 among 153 participants), we also excluded participants whose accuracy in the “math” task was ≤85% correct (20 among 153 participants), since this suggests that they did not attend the very easy math task. Thus, the data of 120 participants (78.4%) are included.

### Stimuli

#### Studies 1-3

The distribution of the stimuli in each of the three experiments was log-uniform (schematically illustrated in Fig. 7). Two categories of trials, randomly selected with equal probability, were intermixed: trials in which both stimuli were pure tones (composed of one frequency *f*) and trials in which both stimuli were harmonic complex tones, composed of three harmonics (fundamental frequency *f*_0_ and the two first overtones: *f*_1_ = 2 ×*f*_0_ and *f*_2_ = 3 ×*f*_0_) having equal intensities. Thus, approximatively half of the trials were pure tones and half harmonic complex tones. The two tones S_1_ and S_2_ in each trial were always of the same category (pure or complex). Pure tones were sampled log-uniformly from 500Hz to 2000Hz (2 octaves). The frequency interval between S_1_ and S_2_ was sampled log-uniformly from the range 0.4-10.1% (uniformly from 6 to 167 cents). The fundamental frequency *f*_0_ of complex tones was sampled log-uniformly from 500Hz to 1000Hz (1 octave). Thus, the first and second overtones *f*_1_ and *f*_2_ ranged from 1000Hz to 2000Hz and from 1500Hz to 3000Hz, respectively. The frequency interval between S_1_ and S_2_ was sampled identically as for the pure tone trials: log-uniformly from the range 0.4-10.1% (uniformly from 6 to 167 cents). In both cases (pure and complex tone-trials), the difference between the frequencies of the two tones S1 and S2 of each trial is much smaller than between the frequencies of the different trials. Importantly, in Experiment 1, the complex tones were composed of all the three harmonics (*f*_0_, *f*_1_, *f*_2_). In Experiment 2, the complex tones did not include the fundamental *f*_0_. In Experiment 3, the complex tones did not include the first overtone *f*_1_. In each of the three experiments, each participant performed 300 trials, divided into three blocks of 100 trials each. Feedback was provided to the participants after each trial and also a summary of the accuracy of their responses after each block.

**Figure 7.**
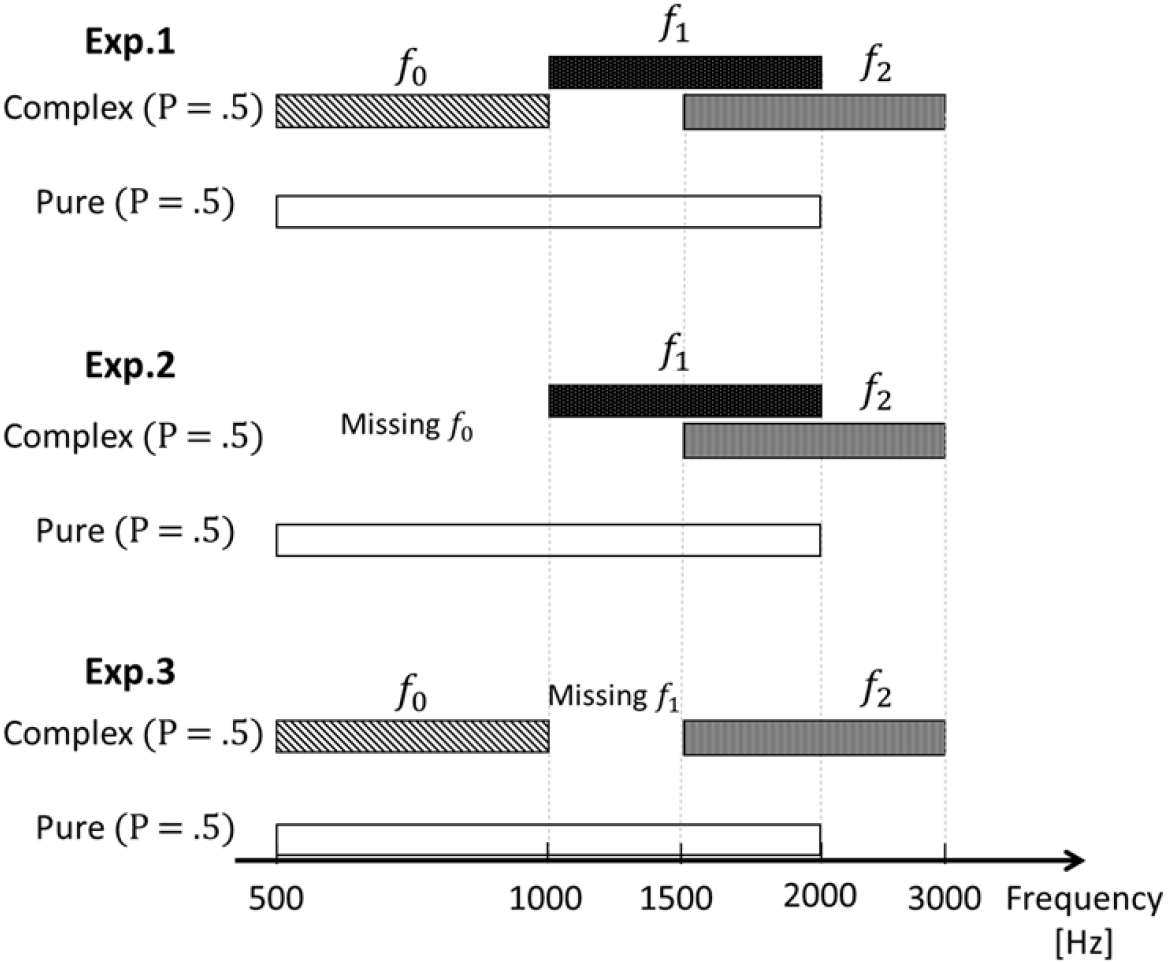
Schematic illustration of stimuli log-uniform distribution in each of the 3 experiments. In all experiments, approximately half of the trials were complex tones and half were pure tones. The two tones composing each trial were always of the same category. Hence, 25% of the sequential trials were pure tone followed by pure tones, 25% pure followed by complex tones, 25% complex followed by complex tones and 25% complex followed by pure tones. In Exp.1 (top) the harmonic complex tones were composed of 3 harmonics – *f*_0_, *f*_1_, *f*_2_. In Exp.2 (mid plot) the complex tones did not include the fundamental frequency *f*_0_. In Exp.3 (bottom), the complex tones did not include the second harmonic *f*_1_. Distributions are shown in binary logarithmic scale.

#### Study 4

All stimuli were pure tones. Stimulus 1 (S_1_) and Stimulus 2 (S_2_) were sampled log-uniformly from 640Hz to 1562Hz (∼1.29 octaves). The frequency interval between S_1_ and S_2_ was sampled log-uniformly from the range 0.4-10.1% (uniformly from 6 to 167 cents). Each participant performed 300 trials, divided into three blocks of 100 trials each. Feedback was provided to the participants after each trial and a summary of the accuracy of their responses after each block.

### Data analysis

Model fitting was performed using the *mgcv* R package^54,55^. Hypothesis tests were performed using the *scikit-learn* and *SciPy* packages in Python. All frequencies are log-transformed.

#### Calculating the contraction bias function by the most recent trial

Following Lieder et al. (2019), we used a Generalized Additive Model (GAM) for estimating the probability that a participant will respond that the second tone 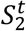 on trial *t*, has a higher pitch.

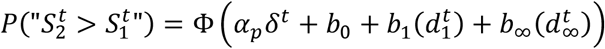

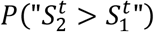 is determined by the standard normal cumulative distribution function Φ (the inverse of the probit link function) of the following sum: frequency difference *δ*^t^ between the (fundamental, in case of complex tones) frequencies of 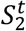 and 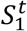, multiplied by the (pre-fitted) participant’s frequency sensitivity *α*_*p*_ (higher *α*_*p*_ means better performance for participant *p*), a constant offset *b*_0_ (representing a constant response preference across all trials and all participants) and the effect of previous trials (*b*_1_ and *b*_∞_). The constant offset *b*_0_ was never found to be significantly different from zero and so is omitted when the model is discussed elsewhere in the text. The effect of previous trials is composed of two non-parametric (spline-estimated) functions *b*_1_ and *b*_∞_, which were shown to be additive^24^: recent (serial dependence) and all other trials.

1. 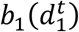 corresponds to the “recent bias” (Lieder et al., 2019), i.e., the contraction to the previous trial, whose magnitude depends on the frequency distance, 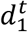, between the frequency of the first tone in the current trial, 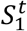, and the frequency of the first tone in the previous trial, 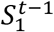. We consider the first tones of consecutive trials because the second tones have close frequencies 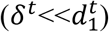 and because the first tone of the previous trial being earlier than the second one, may be more ambiguous. All the predictions of 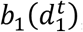, aggregated over all participants, are accompanied by the corresponding standard errors, based on the posterior distribution of the model coefficient 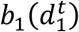.
2. 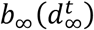 corresponds to the “longer-term effect”^24^ – the contraction to the global mean frequency across the experiment. This term captures the bias toward probable stimuli^56–60^. Its magnitude varies as a function of the frequency distance, 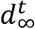, between the frequency of the first tone in the current trial and the mean of the frequencies of the first tones across all trials. Intuitively, 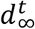 reflects the relative position of the current stimuli within the distribution.

In this paper we are interested in the “recent bias” function, and will not address the longer-term effect. However, we introduce it to our model since it was shown that both biases significantly contribute to the model fit^24^. Importantly, this model has the advantage of separating the effect of recent history from the central tendency induced by the overall average of all previous trials.

#### Fitting single participants (random effects)

In the regression analysis described above, we assumed shared bias functions across participants. This assumption is a statistical necessity since we do not have sufficient individual data to estimate the model individually. To fit single participants, we assumed a shared parameter across participants but allowed small individual deviations (termed *random effects*). Specifically, we added two random effect terms, factorized by individual participants:

1. A random effect on the bias intercept, namely a possible individual response preference bias *b*_0_ that reflects a systematic tendency of a given participant to prefer 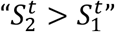 or vice versa regardless of the stimuli.
2. A random effect on the bias slope, that permits comparing the bias magnitudes between individuals, by tuning the bias function. Specifically, we used a metric quantifying the influence of each bias fit – the mean of absolute value of the participant’s bias magnitude: 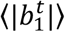, where 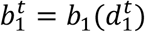 is the bias magnitude in trial *t*.

These random effect terms allowed us to compare serial dependence magnitudes quantitatively, and account for inter-individual variability. Because assumed deviations are small, inference in these models is tractable.

#### Varying-coefficient models and lapse rate

A single model was fit for all participants, with common bias functions, but with participant specific sensitivities *α*_*p*_. We combined the pre-fitted *α*-participant pairing in the model using the varying-coefficients models method^61^. This method assumes linearity in the regressors, but their coefficients are allowed to change smoothly with the value of other variable (alphas in our case): 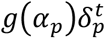. Here *α*_*p*_ is the pre-fitted sensitivity parameter, 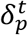 is the difference between the target tones for each participant *p*, and *g* is some smooth fitted function. We added a lapse parameter *λ* to account for occasional inattentiveness^62^: 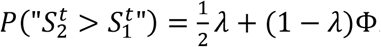. We set *λ* to a fixed 0.05.

#### Study 1: Comparison between the four types of trials on aggregated data

The aim of this analysis was to characterize and compare the recent bias in pitch arising in the four types of consecutive trials (pure after pure, complex after complex, pure after complex and complex after pure). For this purpose, we aggregated the data of the 3 experiments. For fair comparison, we considered only preceding pure-tone trials (recent bias inducers) that were in the same frequency range (500-1000Hz, lower octave) as the fundamental frequencies in complex tones trials (Methods: *Stimuli*). For each type of trials, we calculated the recent bias (in bold) using GAM as follows:

- Complex after complex:

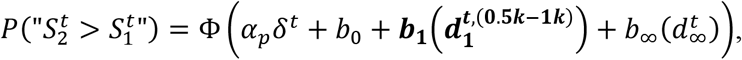

where 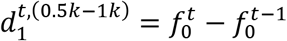 is the frequency distance between the fundamental of the first tone in the current and the fundamental of the first tone in the previous trials. 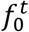 and 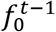 are both in the range 500-1000Hz. A total of 58,779 trials were included.
- Complex after pure (lower octave)

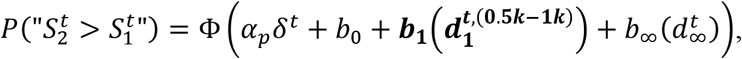

where 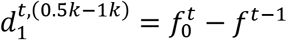 is the frequency distance between the fundamental of the first tone in the current trial and the first pure tone of the previous trial. 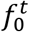 and *f*^*t*−1^ are both in the range 500-1000Hz. As we considered only pure tones in the lower octave (half of the trials), a total of 29,577 trials were included.
- Pure (both octaves) after pure (lower octave):

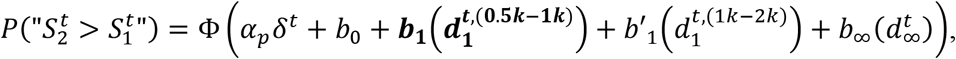

where 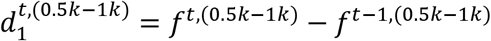 is the frequency distance when the first pure tone of the current trial is in the lower octave range (same octave as the first pure tone of the preceding trial). 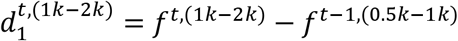 is the frequency distance when the first pure tone of the current trial is in the upper octave range (higher octave than the first pure tone of the previous trial). Since we considered only pure tones in the lower octave at *t*−1 (half of the data), a total of 29,515 trials were included.
- Pure (both octaves) after complex:

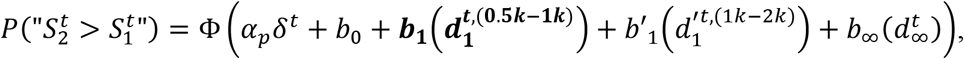

where 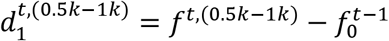 is the frequency distance when the first pure tone of the current trial is in the lower octave range (same octave as the fundamental of the previous trial). 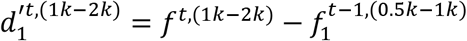 is the frequency distance between the first pure tone of the current trial (ranging in the upper octave range) and the first tone’s second harmonic of the previous trial (ranging in the same upper octave). A total of 59,027 trials were included.

#### Studies 2 and 3: Contraction at the level of frequency channels

In these analyses, we focused on consecutive trials from different categories. The aim was to measure the contraction at the level of specific harmonics. We used the same model as in Study 1 except that, for simplicity, we used linear regressions, as they performed as well as spline functions in these cases. Indeed, the effect measured across-categories was substantially weaker and noisier at larger frequency distance. We believe that more data will yield similar curves as those obtained within categories, and will improve the fit of the function.

Thus, for pure (both octaves) after complex trials, the model takes the following form:

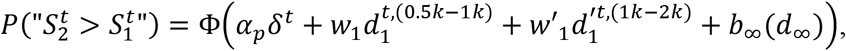

where 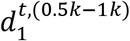 is the frequency distance between the first pure tone in the lower octave and the fundamental frequency of the first complex tone;

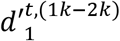 is the frequency distance between the pure tone in the upper octave and the second harmonic of the first complex tone;

*w*_1_ and 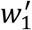 are the corresponding slopes of the recent bias.

There are two components for the recent bias, describing the contraction by the fundamental frequency *f*_0_ of the pure tone is in the lower octave (500-1000Hz) (Study 2, Pure after Complex) and the contraction by the second harmonic *f*_1_ of the pure tone is in the upper octave (1000-2000Hz) (Study 3).

Similarly, for complex after pure (lower octave) trials, the model takes the following form, using the same notation:

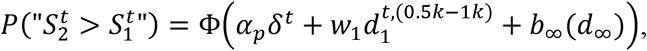

where there is one component for the recent bias, describing the contraction by the pure tone (*f*) of the fundamental frequency (*f*_0_) when the latter is in the lower octave (Study 2, Complex after Pure).

## Acknowledgements

This project has received funding from the European Research Council (ERC) under the European Union’s Horizon 2020 research and innovation program (grant agreement No 833694) and the Israel Science Foundation (Grant No. 1650/17), both awarded to Merav Ahissar.

## Authors’ Contributions

Initialization of the project: I.L. Design of the experiments: I.L. and M.A. Conceptualization of the study: I.L., A.S., M.A. Collection of data: I.L. Analysis of the data: I.L, A.S. Funding acquisition and supervision: M.A. All authors contributed to the interpretation of data and writing of the manuscript.

These authors contributed equally: Itay Lieder, Aviel Sulem.

## Material availability statement

The two-tone discrimination experiments were conducted online via the Amazon Mechanical Turk crowd-sourcing platform. The experiments were conducted in JavaScript and administered using web browsers with tones that were preloaded and played using HTML5 Audio. We used Psychtoolbox-3 MATLAB toolbox (http://psychtoolbox.org/) for creating the auditory stimuli. Analysis was conducted using the ‘mixed GAM computation vehicle with automated smoothness estimation’ (mgvc) free package https://cran.r-project.org/web/packages/mgcv/index.html. The data sets generated and analyzed during the present studies are available at https://github.com/avielsulem/PitchSerialEffects.git.

## References

1. Ahissar, M. & Hochstein, S. The reverse hierarchy theory of visual perceptual learning. Trends Cogn Sci 8, 457–464 (2004).

2. de Lange, F. P., Heilbron, M. & Kok, P. How Do Expectations Shape Perception? Trends Cogn Sci 22, 764–779 (2018).

3. Seriès, P. & Seitz, A. R. Learning what to expect (in visual perception). Frontiers in Human Neuroscience Preprint at https://doi.org/10.3389/fnhum.2013.00668 (2013).

4. Klink, P. C., van Wezel, R. J. A. & van Ee, R. United we sense, divided we fail: Context-driven perception of ambiguous visual stimuli. Philosophical Transactions of the Royal Society B: Biological Sciences vol. 367 932–941 Preprint at https://doi.org/10.1098/rstb.2011.0358 (2012).

5. Raviv, O., Lieder, I., Loewenstein, Y. & Ahissar, M. Contradictory Behavioral Biases Result from the Influence of Past Stimuli on Perception. PLoS Comput Biol 10, (2014).

6. Snyder, J. S., Schwiedrzik, C. M., Vitela, A. D. & Melloni, L. How previous experience shapes perception in different sensory modalities. Front Hum Neurosci 9, (2015).

7. Fritsche, M., Mostert, P. & de Lange, F. P. Opposite Effects of Recent History on Perception and Decision. Current Biology 27, 590–595 (2017).

8. Fründ, I., Wichmann, F. A. & Macke, J. H. Quantifying the effect of intertrial dependence on perceptual decisions. J Vis 14, 1–16 (2014).

9. Cicchini, G. M., Mikellidou, K. & Burr, D. Serial dependencies act directly on perception. J Vis 17, (2017).

10. Fischer, J. & Whitney, D. Serial dependence in visual perception. Nat Neurosci 17, 738–743 (2014).

11. Manassi, M., Liberman, A., Chaney, W. & Whitney, D. The perceived stability of scenes: Serial dependence in ensemble representations. Sci Rep 7, (2017).

12. Bliss, D. P., Sun, J. J. & D’Esposito, M. Serial dependence is absent at the time of perception but increases in visual working memory. Sci Rep 7, (2017).

13. Manassi, M., Liberman, A., Kosovicheva, A., Zhang, K. & Whitney, D. Serial dependence in position occurs at the time of perception. Psychon Bull Rev 25, 2245– 2253 (2018).

14. Liberman, A., Fischer, J. & Whitney, D. Serial dependence in the perception of faces. Current Biology 24, 2569–2574 (2014).

15. Turbett, K., Palermo, R., Bell, J., Hanran-Smith, D. A. & Jeffery, L. Serial dependence of facial identity reflects high-level face coding. Vision Res 182, 9–19 (2021).

16. Liberman, A., Manassi, M. & Whitney, D. Serial dependence promotes the stability of perceived emotional expression depending on face similarity. Atten Percept Psychophys 80, 1461–1473 (2018).

17. Taubert, J., Alais, D. & Burr, D. Different coding strategies for the perception of stable and changeable facial attributes. Sci Rep 6, (2016).

18. Xia, Y., Leib, A. Y. & Whitney, D. Serial dependence in the perception of attractiveness. J Vis 16, (2016).

19. van der Burg, E., Toet, A., Brouwer, A. M. & van Erp, J. B. F. Sequential Effects in Odor Perception. Chemosens Percept 15, 19–25 (2022).

20. Arzounian, D., de Kerangal, M. & de Cheveigné, A. Sequential dependencies in pitch judgments. J Acoust Soc Am 142, 3047–3057 (2017).

21. Chambers, C. et al. Prior context in audition informs binding and shapes simple features. Nat Commun 8, (2017).

22. Chambers, C. & Pressnitzer, D. Perceptual hysteresis in the judgment of auditory pitch shift. Atten Percept Psychophys 76, 1271–1279 (2014).

23. Motala, A., Zhang, H. & Alais, D. Auditory Rate Perception Displays a Positive Serial Dependence. Iperception 11, (2020).

24. Lieder, I. et al. Perceptual bias reveals slow-updating in autism and fast-forgetting in dyslexia. Nat Neurosci 22, 256–264 (2019).

25. Liberman, A., Zhang, K. & Whitney, D. Serial dependence promotes object stability during occlusion. J Vis 16, (2016).

26. Pascucci, D. et al. Laws of concatenated perception: Vision goes for novelty, decisions for perseverance. PLoS Biol 17, (2019).

27. Papadimitriou, C., Ferdoash, A. & Snyder, L. H. Ghosts in the machine: memory interference from the previous trial. J Neurophysiol 113, 567–577 (2015).

28. Papadimitriou, C., White, R. L. & Snyder, L. H. Ghosts in the Machine II: Neural correlates of memory interference from the previous trial. Cerebral Cortex 27, 2513– 2527 (2017).

29. Mei, G., Chen, S. & Dong, B. Working memory maintenance modulates serial dependence effects of perceived emotional expression. Front Psychol 10, (2019).

30. Ceylan, G., Herzog, M. H. & Pascucci, D. Serial dependence does not originate from low-level visual processing. Cognition 212, (2021).

31. st. John-Saaltink, E., Kok, P., Lau, H. C. & de Lange, F. P. Serial dependence in perceptual decisions is reflected in activity patterns in primary visual cortex. Journal of Neuroscience 36, 6186–6192 (2016).

32. Hochstein, S. & Ahissar, M. View from the Top: Hierarchies and Reverse Hierarchies in the Visual System. Neuron 36, 791–804 (2002).

33. Cicchini, G. M., Benedetto, A. & Burr, D. C. Perceptual history propagates down to early levels of sensory analysis. Current Biology 31, 1245-1250.e2 (2021).

34. Ahissar, M., Nahum, M., Nelken, I. & Hochstein, S. Reverse hierarchies and sensory learning. Philosophical Transactions of the Royal Society B: Biological Sciences vol. 364 285–299 Preprint at https://doi.org/10.1098/rstb.2008.0253 (2009).

35. Nelken, I. & Ahissar, M. High-level and low-level processing in the auditory system: the role of primary auditory cortex. in Dynamics of speech production and perception vol. 374 343–353 (2006).

36. Chialvo, D. R. How we hear what is not there: A neural mechanism for the missing fundamental illusion. Chaos 13, 1226–1230 (2003).

37. de Cheveigné, A. Pitch perception. The Oxford Handbook of Auditory Science: Hearing (Oxford University Press, 2010). doi:10.1093/oxfordhb/9780199233557.013.0004.

38. Zatorre, R. J. Finding the missing fundamental. Nature 436, 1093–1094 (2005).

39. Oxenham, A. J. Pitch perception. Journal of Neuroscience vol. 32 13335–13338 Preprint at https://doi.org/10.1523/JNEUROSCI.3815-12.2012 (2012).

40. Plomp, R. Pitch of Complex Tones. J Acoust Soc Am 41, 1526–1533 (1967).

41. Moore, B. C. J. An Introduction to the Psychology of Hearing. (Emerald, 2012).

42. Näätänen, R., Paavilainen, P., Rinne, T. & Alho, K. The mismatch negativity (MMN) in basic research of central auditory processing: A review. Clinical Neurophysiology vol. 118 2544–2590 Preprint at https://doi.org/10.1016/j.clinph.2007.04.026 (2007).

43. Chen, Y., Huang, X., Luo, Y., Peng, C. & Liu, C. Differences in the neural basis of automatic auditory and visual time perception: ERP evidence from an across-modal delayed response oddball task. Brain Res 1325, 100–111 (2010).

44. van der Burg, E., Rhode, G. & Alais, D. Positive sequential dependency for face attractiveness perception. J Vis 19, 1–16 (2019).

45. Kiyonaga, A., Scimeca, J. M., Bliss, D. P. & Whitney, D. Serial Dependence across Perception, Attention, and Memory. Trends Cogn Sci 21, 493–497 (2017).

46. Fritsche, M. & de Lange, F. P. The role of feature-based attention in visual serial dependence. J Vis 19, (2019).

47. Fornaciai, M. & Park, J. Attractive Serial Dependence in the Absence of an Explicit Task. Psychol Sci 29, 437–446 (2018).

48. Kathmann, N., Frodl-Bauch, T. & Hegerl, U. Stability of the mismatch negativity under different stimulus and attention conditions. (1999).

49. Manassi, M., Kristjánsson, Á. & Whitney, D. Serial dependence in a simulated clinical visual search task. Sci Rep 9, (2019).

50. Manassi, M. & Whitney, D. Illusion of visual stability through active perceptual serial dependence. vol. 8 https://www.science.org (2022).

51. Fiser, J., Berkes, P., Orbán, G. & Lengyel, M. Statistically optimal perception and learning: from behavior to neural representations. Trends Cogn Sci 14, 119–130 (2010).

52. Jaffe-Dax, S., Frenkel, O. & Ahissar, M. Dyslexics’ faster decay of implicit memory for sounds and words is manifested in their shorter neural adaptation. Elife 6, (2017).

53. Lu, Z. L., Williamson, S. J. & Kaufman, L. Behavioral lifetime of human auditory sensory memory predicted by physiological measures. Science (1979) 258, 1668–1670 (1992).

54. Wood, S. Package mgcv. http://cran.r-project.org/web/packages/mgcv/mgcv.pdf.

55. Knoblauch, K. CRAN - Package psyphy: Functions for Analyzing Psychophysical Data in R. CRAN https://cran.r-project.org/web/packages/psyphy/index.html (2022).

56. Ashourian, P. & Loewenstein, Y. Bayesian inference underlies the contraction bias in delayed comparison tasks. PLoS One 6, (2011).

57. Hollingworth, H. L. The Central Tendency of Judgment. Source: The Journal of Philosophy vol. 7 https://about.jstor.org/terms (1910).

58. Jaffe-Dax, S., Lieder, I., Biron, T. & Ahissar, M. Dyslexics’ usage of visual priors is impaired. J Vis 16, 1–9 (2016).

59. Karim, M., Harris, J. A., Langdon, A. & Breakspear, M. The influence of prior experience and expected timing on vibrotactile discrimination. Front Neurosci (2013) doi:10.3389/fnins.2013.00255.

60. Olkkonen, M., McCarthy, P. F. & Allred, S. R. The central tendency bias in color perception: Effects of internal and external noise. J Vis 14, (2014).

61. Hastie, T. & Tibshirani, R. Varying-Coefficient Models. Source: Journal of the Royal Statistical Society. Series B (Methodological) vol. 55 (1993).

62. Dai, H. & Micheyl, C. Psychometric functions for pure-tone frequency discrimination. J Acoust Soc Am 130, 263–272 (2011).

